# Development and decline of the hippocampal long-axis specialization and differentiation during encoding and retrieval of episodic memories

**DOI:** 10.1101/323097

**Authors:** Espen Langnes, Didac Vidal-Piñeiro, Markus H. Sneve, Inge K. Amlien, Kristine B Walhovd, Anders M Fjell

**Author notes:** Address correspondence to: Anders M Fjell, Pb. 1094 Blindern, 0317 Oslo, Norway.

## Abstract

Change in hippocampal function is a major factor in lifespan development and decline of episodic memory. Evidence indicates a long-axis specialization where anterior hippocampus is more engaged during encoding and posterior during retrieval. We tested the lifespan trajectory of hippocampal long-axis episodic memory-related activity and functional connectivity (FC). 496 participants (6.8-80.8 years) were scanned with functional MRI while encoding and retrieving associative memories. We found clear evidence for a long-axis encoding-retrieval specialization. These long-axis effects declined linearly during development and aging, eventually vanishing in the older adults. This was mainly driven by age effects on retrieval. Retrieval was associated with gradually lower activity from childhood to adulthood, followed by positive age-relationships until 70 years. Interestingly, this pattern characterized task engagement regardless of memory success or failure. Children engaged posterior hippocampus more than anterior, while anterior hippocampus was more activated relative to posterior already in teenagers. Intra-hippocampal connectivity increased during task, and this increase declined with age. In sum, the results suggest that hippocampal long-axis differentiation and communication during episodic memory tasks develop rapidly during childhood and adolescence, are markedly different in older compared to younger adults, and are related to task engagement, not the successful completion of the task.

## Introduction

Episodic memory function declines in normal aging (Nyberg et al.; Ronnlund et al. 2005), with changes in hippocampal function appearing to be a major cause (Fjell et al. 2014). However, although hippocampus (HC) is critical for encoding, consolidation and retrieval of episodic memories (Schacter et al. 2012; Tulving 1984, 2002), its specific role is still debated (Moscovitch et al. 2016; Tulving 2002, 1984; Schacter et al. 2012; Klein 2014). What is clear is that the role of HC during encoding and retrieval is not uniform, and that the different aspects of hippocampal involvement must be supported by partly different anatomical regions or subfields closely communicating to allow successful episodic memory operations (Moscovitch et al. 2016; Strange et al. 2014; Collin, Milivojevic, and Doeller 2015). For instance, recent studies have suggested a long axis specialization of HC in memory processing (Poppenk et al. 2013; Chase et al. 2015; Kühn and Gallinat 2014), where the anterior hippocampus (aHC) is especially engaged in encoding, and the posterior hippocampus (pHC) is more heavily engaged in retrieval and reconstruction (Kühn and Gallinat 2014; Lepage, Habib, and Tulving 1998; Poppenk et al. 2013; Nadel, Hoscheidt, and Ryan 2012). The purpose of the present study is to test whether hippocampal activity across the anterior-posterior long axis during encoding and retrieval relates to chronological age in development, adulthood and aging, and whether such differences in long axis specialization are specifically related to successful encoding or retrieval of episodic memories. To this end, 496 cognitively healthy participants from 6.8 to 80.8 years underwent functional magnetic resonance imaging (fMRI) during encoding and retrieval in an associative memory task, allowing testing of activity and connectivity differences in and between the aHC vs pHC.

Previous studies have shown age-effects on activation differences between aHC and pHC. It has been suggested that protracted structural and functional development of hippocampal sub-regions along the anterior-posterior axis contributes to age-related differences in episodic memory performance in children and youth (DeMaster et al. 2014; DeMaster and Ghetti 2013; Sastre Iii et al. 2016). Towards the other end of the lifespan, the literature is divergent. While many studies find age-related reductions in hippocampal activity associated with successful episodic memory (Cabeza et al. 2004; Daselaar et al. 2006; Dennis et al. 2008; Dennis, Kim, and Cabeza 2008; Murty et al. 2009), others find lack of age effects (Cansino et al. 2015; de Chastelaine et al. 2016b; Duverne, Habibi, and Rugg 2008; Park et al. 2013) (for reviews, see (Leal and Yassa 2013; Nyberg 2017), or that differences are affected by factors such as task performance (de Chastelaine et al. 2016a). Longitudinal studies show that hippocampal activity may be preserved in older adults with stable memory (Pudas et al. 2013) and reduced in those who decline (Persson et al. 2012). Recent results also suggest that age-correlations may be restricted to certain hippocampal regions (Carr et al. 2017), such as aHC during encoding (Salami, Eriksson, and Nyberg 2012; Daselaar et al. 2003). Thus, it is possible that selective age effects on hippocampal long-axis specialization exist both in development and in aging. Hence, a large-scale investigation of long-axis activity during both encoding and retrieval through the lifespan is needed.

Although any functional specialization will require close communication between aHC and pHC, tracer studies in animals have revealed few direct connections. Rather, different parts of the hippocampus seem to display distinctive, topographically arranged, neuronal connectivity patterns (Fanselow and Dong 2010). Still, aHC and pHC have multiple routes through which they interact to ensure coordinated information processing (Fanselow and Dong 2010), and a reasonable hypothesis would thus be that the communication between them increases during memory processing (Robinson, Salibi, and Deshpande 2016). A recent meta-analysis found partially overlapping and partly separate connectivity patterns between aHC vs. pHC and the rest of the cortex using task-related fMRI as well as diffusion tensor imaging (Robinson, Salibi, and Deshpande 2016). Studies focusing on successful episodic memory retrieval typically (Ranganath et al. 2005; Schott et al. 2013) find increased connectivity between the hippocampus and other cortical areas (but see (King et al. 2015)). Source memory-related connectivity has been reported to be higher in younger than older (King, de Chastelaine, and Rugg 2017) but to our knowledge, age effects on intra-hippocampal connectivity-changes have not been tested.

### Hypotheses

The main aim of the study is to test degree of hippocampal long-axis specialization and differentiation during encoding and retrieval through the lifespan, with regard to activity and connectivity. With specialization, we refer to a two-way long-axis (anterior vs. posterior HC) × condition (encoding vs. retrieval) interaction. Of similar interest is what we refer to as age differentiation, which represents an effect of age on long-axis differences within each of the conditions, i.e. encoding and retrieval. In the present study, we tested both HC specialization as well as age effects on HC differentiation.

We hypothesized that:

1. An encoding-retrieval hippocampal long-axis specialization will be seen, in that aHC will be more activated during encoding than pHC, and pHC will be more activated during retrieval than aHC.
2. The long-axis functional differentiation will show an inverted U-shape through the lifespan, i.e. be more evident with increasing age in development, reach a plateau in young adults, and then break down in older adults.
3. FC between aHC and pHC will increase during both encoding and retrieval.
4. Task-related FC increases will overall show a U-shaped age-trajectory, with higher FC in children and possibly older adults compared to younger adults. This hypothesis was speculative, based on indirect evidence from previous studies suggesting that children (Sastre Iii et al. 2016) and older adults (Salami, Pudas, and Nyberg 2014) tend to use the hippocampus in a less specialized way, possibly due to neural inefficiency or lack of inhibition.

## Materials and Methods

### Participants

The participants were recruited from ongoing studies coordinated by the Center for Lifespan Changes in Brain and Cognition (LCBC) at the Department of Psychology, University of Oslo, Norway. The final sample consisted of 496 well-screened cognitively healthy participants (337 females, age, 6.8-80.8 years; mean 39.1 years, standard deviation = 17.6 years). All participants gave written informed consent, and the Regional Ethical Committee of South Norway approved the study. The participants reported no history of neurological or psychiatric disorders, chronic illness, premature birth, learning disabilities, or use of medicines known to affect nervous system functioning. At scanning a separate clinical sequence (T2-FLAIR) was included for neurological evaluation by a neuroradiologist, and the scans were required to be free of significant injuries or conditions They were further required to speak fluent Norwegian, and have normal or corrected-to-normal hearing and vision. The participants were compensated for their participation. The participants were required to score ≥26 on the Mini Mental State Examination (Folstein, Folstein, and McHugh 1975). Participants above 8 years were tested on Vocabulary and Matrix Reasoning subtests of the Wechsler’s Abbreviated Scale Intelligence Scale (WASI) (Wechsler, 1999). Participants under the age of eight years were tested on the same subtest from the Wechsler Preschool and Primary Scale of Intelligence (WIPPSI-III) (Wechsler 2002). All scored within the normal IQ range (>85) and a T-score of ≤30 on the California Verbal Learning Test II— Alternative Version (CVLTII) (Delis et al. 2008) immediate delay and long delay. Participants were further excluded due to experimental and operator errors (incorrect order of the sequence, participants failing to understand the task, disabled button response, etc.), low number of trials available for fMRI analysis (n = 24; <6 per condition of interest) and extreme movement (n = 1; >1.5 mm mean movement). Participant demographics are summarized in Table 1. The sample partially overlaps with the samples used in Sneve et al. (Sneve et al. 2015), where encoding activity for 78 adults were included, and Vidal-Piñero et al. (Vidal-Piñeiro et al. 2017), where encoding and retrieval activity for 143 adult participants were analysed. In neither of these studies were activity along the hippocampal long axis studied.

**Table 1.** Main demographic and neuropsychological variables for the complete sample, as well as broken down in 8 separate age groups. ^a^n = 410; ^b^n = 352; ^c^n = 319; ^d^n = 459; ^e^n = 457.

| Group | All | Age1 | Age2 | Age3 | Age4 | Age5 | Age6 | Age7 | Age8 |
| --- | --- | --- | --- | --- | --- | --- | --- | --- | --- |
| n | 496 | 15 | 30 | 161 | 90 | 59 | 52 | 60 | 28 |
| Females/males | 337:158 | 10:5 | 12:18 | 121:40 | 66:24 | 42:17 | 37:15 | 36:24 | 12:16 |
| Age range | 6.8-80.8 | 6.8-13.9 | 14.4-19.6 | 20.1-29.9 | 30.0-39.9 | 40.1-50.0 | 50.0-60.0 | 60.0-69.9 | 70.0-80.8 |
| Age mean (SD) | 39.1 (17.6) | 9.9 (2.2) | 16.9 (1.2) | 25.6 (2.7) | 33.9 (2.7) | 45.6 (2.7) | 55.1 (2.9) | 65.0 (3.1) | 74.0 (3.4) |
| MMSE<sup>a</sup> | 29.1 (1.0) | -- | -- | 29.2 (1.0) | 29.4 (0.8) | 29.2 (1.0) | 29.0 (0.9) | 28.7 (1.2) | 28.3 (1.4) |
| CFT Recall<sup>b</sup> | 21.0 (6.4) | 19.1 (6.6) | 30.6 (3.7) | 22.5 (5.7) | 21.2 (6.3) | 22.3 (5.6) | 19.1 (6.4) | 17.8 (6.1) | 17.3 (5.8) |
| Vocabulary<sup>c</sup> | 62.3 (9.1) | 37.7 (12.7) | 56.1 (4.9) | 62.6 (6.1) | 64.1 (6.4) | 63.4 (8.0) | 65.9 (5.0) | 65.3 (7.5) | 66.8 (7.4) |
| Matrices<sup>d</sup> | 28.3 (6.1) | 23.7 (5.5) | 29.9 (2.8) | 29.0 (3.7) | 29.3 (2.7) | 28.9 (6.5) | 28.7 (9.7) | 25.6 (5.0) | 25.7 (13.0) |
| CVLT Learning<sup>e</sup> | 58.4 (9.5) | 50.6 (11.1) | 61.2 (7.9) | 61.5 (8.3) | 61.0 (8.6) | 59.3 (9.4) | 55.6 (8.9) | 52.6 (9.6) | 51.4 (8.5) |
| CVLT 30'<sup>e</sup> | 13.0 (2.5) | 10.9 (2.9) | 13.4 (2.0) | 13.7 (2.2) | 13.9 (2.1) | 13.3 (2.3) | 12.4 (2.3) | 11.6 (2.9) | 10.9 (2.7) |

### Experimental design - fMRI tasks

Participants were scanned using BOLD fMRI during an experimental task that consisted of an incidental encoding task and a subsequent memory test after ≈ 90 minutes. The memory task was optimized to allow for the investigation of individual differences in item-source associative memory performance, i.e., the ability to remember a previously encountered item together with information about the encoding context. This task was optimized to allow us to investigate the neural correlates of source memory/ associative memory. A schematic presentation of the design is shown in Figure 1, and the task has also been described in Sneve et al. (Sneve et al. 2015). The participants were verbally instructed minutes before both experimental tasks and did not go through any practice session before entering the scanner.

**Figure 1.**
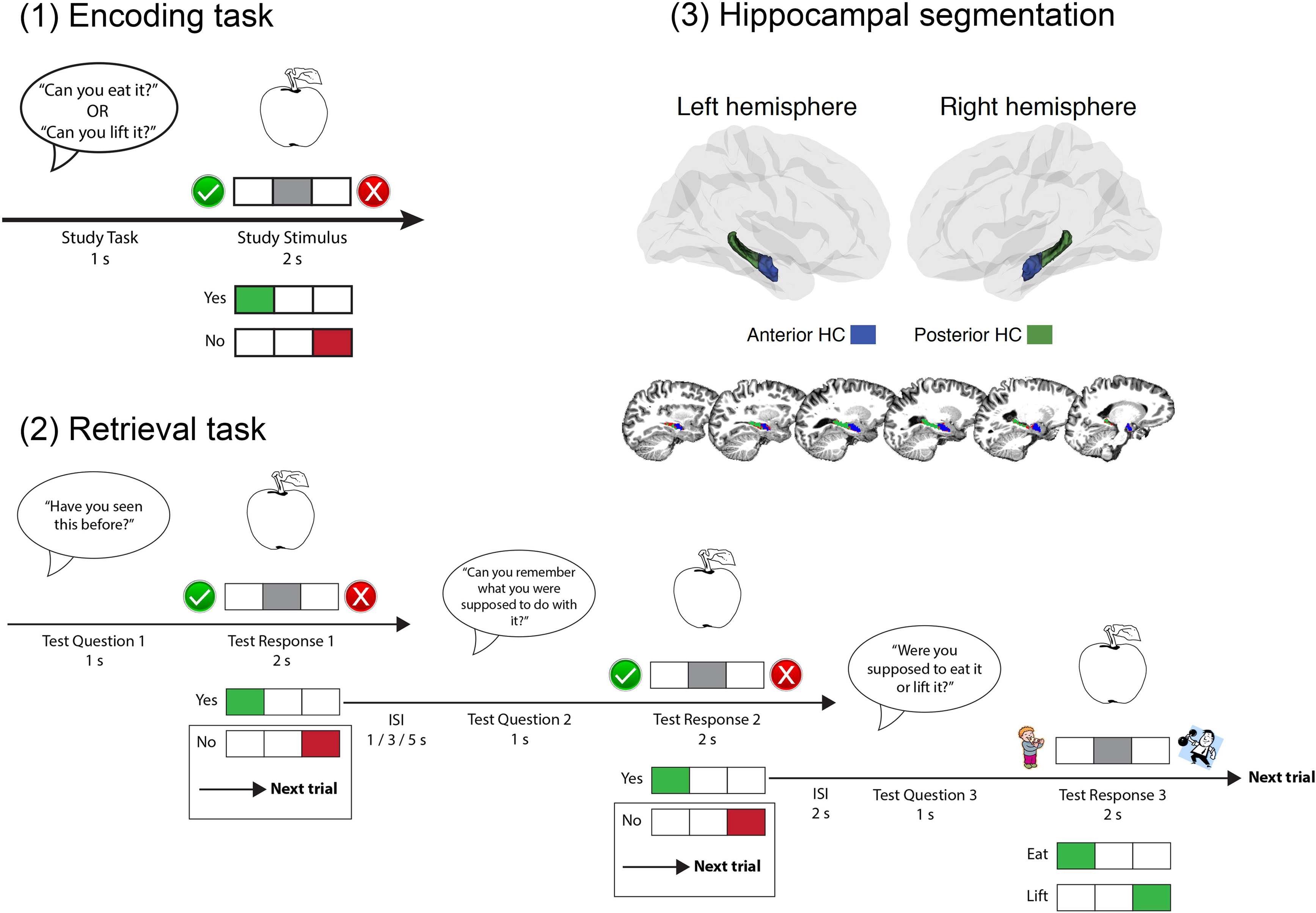
Experimental paradigm - fMRI task. (1) Schematic overview of an encoding trial. The green (?) and the red (X) symbols were present on the screen to indicate which button indicated “Yes” and “No”, respectively. (2) Schematic overview of the test condition of the experiment Test Questions 1 and 2 required a Yes/No response whereas Question 3 consisted of a two-alternative forced choice task. The trial ended if the participant responded “No” to either one of the two first questions. ISI: InterStimulus Interval, s: second. Adapted from Sneve et al. (Sneve et al. 2015). (3) Overview of the hippocampal long-axis segmentation scheme. Top panel: aHC and pHC in MNI305-space. Bottom panel: sagittal view of the functional definitions of left aHC and pHC for a representative participant in native subject space. Voxels colored red represent the FreeSurfer-segmented left hippocampus in high-resolution (1mm^3^) structural space. AHC (blue) / pHC (red) voxels are overlaid in functional resolution (27mm^3^).

The encoding and the retrieval tasks consisted of two and four runs, respectively, that included 50 trials each. All runs started and ended with a 11s baseline period, which was also presented once in the middle of each run. The stimulus material consisted of 300 black and white line drawings depicting everyday objects and items. During encoding, the participants went through 100 trials of a task in which they performed simple evaluations of everyday objects and items. A trial had the following structure: a female voice asked either “Can you eat it?” or “Can you lift it?” Both questions were asked equally often and were pseudorandomly mixed across the different objects. One second after question onset, a black and white line drawing of an object was presented on the screen along with response indicators. Participants were instructed to produce yes/no-responses based on their subjective evaluations of object/task-contingencies, and that there were no correct responses to the task. Button response was counterbalanced across participants. The object remained in the screen for 2 s, when it was replaced by a central fixation cross that remained throughout the intertrial interval (ITI; 1-7s exponential distribution over four discrete intervals).

During the surprise memory test, 200 line drawings of objects were presented; 100 of these had been shown and evaluated during encoding while the remaining 100 objects were new. A test trial started with the presentation of an object (old or new, pseudorandomly picked) and the question (Question 1) “Have you seen this item before?”. Each object stayed on the screen for 2 seconds. Participants were instructed to respond “Yes” if they remembered seeing the item during the encoding condition, and “No” otherwise. If the participant indicated that (s)he remembered seeing the object, a new question followed (Question 2): “Can you remember what you were asked to do with the item?” A “Yes”-response to this question, indicating that the participant also remembered the action associated with the object during encoding, led to a final two-alternative forced choice question (Question 3): “Were you asked to eat it or lift it?” Here, the participant indicated either “Eat” ("I evaluated whether I could eat the item during the encoding condition)" or “Lift” ("I evaluated lifting the item"). The specific questions asked during scanning were simplified to fit within the temporal limits of the paradigm. Despite the response-dependent nature of the fMRI regressors, the design efficiency was tentatively optimized to ensure sufficient complexity in the recorded time series (http://surfer.nmr.mgh.harvard.edu/optseq/).

### Analysis of behavioural data

The main behavioral measure of interest was the source memory score. A participant’s raw source memory score was calculated as the proportion of encoded items that were recognized with correct source memory of the associated encoding action. Source memory for a trial was considered when: a participant correctly recognized an item (correct “Yes” response to test Question 1), stated that (s)he remembered the associated action (“Yes” response to test Question 2), and picked the correct associated action in the two-alternative forced-choice question (correct response to Question 3). A corrected source memory score was calculated from the raw source memory score by subtracting the number of times a participant produced a wrong source response (i.e., wrong response to test Question 3). This correction tentatively accounts for processes such as false memories, threshold criteria in Question 2 or guessing (Vidal-Pineiro et al. 2017). Corrected source memory scores was the only behaviour measure considered in the analyses.

### MR1 scanning and preprocessing

Imaging was performed at a Siemens Skyra 3T MRI unit with a 24-channel head coil at Rikshospitalet, Oslo University Hospital. For the functional imaging scanning the parameters were equivalent across all runs: 43 slices (transversal, no gap) were measured using T2* weighted BOLD EPI (TR=2390ms; TE=30ms; flip angle=90°; voxel size=3x3x3mm; FOV=224x224; interleaved acquisition; GRAPPA=2). Each encoding run produced 131 volumes while the number of volumes per retrieval run was dependent on participants’ responses (mean 207 volumes). Three dummy volumes were collected at the start of each fMRI run to avoid T1 saturation effects in the analyzed data. Additionally, a standard double-echo gradient-echo field map sequence was acquired for distortion correction of the EPI images. Anatomical T1-weighted MPRAGE images consisting of 176 sagittally oriented slices were obtained using a turbo field echo pulse sequence (TR = 2300 msec, TE = 2.98 msec, flip angle = 8°, voxel size = 1 × 1 × 1 mm, FOV= 256 × 256 mm). Visual stimuli were presented in the scanner environment with a 32-inch InroomViewing Device monitor while participants responded using the ResponseGrip device (both NordicNeuroLab, Norway). Auditory stimuli were presented to the participants’ headphones through the scanner intercom.

Cortical reconstruction and volumetric segmentation of the T1-weighted scans were performed with FreeSurfer 5.3 https://surfer.nmr.mgh.harvard.edu/ This processing included segmentation of the subcortical white matter and deep grey matter volumetric structures (including the hippocampus) (Fischl et al., 2004a, 2002), surface inflation (Fischl et al., 1999a), and registration to a spherical atlas which utilized individual cortical folding patterns to match cortical geometry across subjects (Fischl et al., 1999b).

fMRI data was initially corrected for B0 inhomogeneity, motion and slice timing corrected, smoothed (5mm FWHM) in volume space and high-pass filtered (at 0.01Hz) using FSL (http://fsl.fmrib.ox.ac.uk/fsl/fslwiki). Next, FMRIB’s ICA-based Xnoiseifier (FIX) (Salimi-Khorshidi et al., 2014) was used to auto-classify noise components and remove them from the fMRI data. The classifier was trained on a task-specific dataset in which task fMRI data from 36 participants had been manually classified into signal and noise components (age span in training set: 7-80; fMRI acquisition parameters identical to the current study]. Motion confounds (24 parameters) were regressed out of the data as a part of the FIX routines. Transformation matrices between functional-native, structural-native and freesurfer average space were computed to delineate hippocampal structures and and bring them to the functional-native space. Next, the preprocessed fMRI data, at the functional space, was introduced in a first-level GLM analysis.

### fMRI analysis

A first-level general linear model (GLM) was set up with FSFAST (https://surfer.nmr.mgh.harvard.edu/fswiki/FsFast) for each encoding and retrieval run, consisting of several conditions/regressors modeled as events with onsets and durations corresponding to the trial events during encoding and retrieval and convolved with a two-gamma canonical hemodynamic response function (HRF), At retrieval, each "old" trial (test item presented during encoding, n = 100) was assigned to a condition based on the participant’s response at test. Two conditions of interest were modeled both at encoding and at retrieval. 1) The source memory encoding condition consisted of items that were later correctly recognized with correct source memory (Yes response to test Questions 1 and 2 and correct response to Question 3). 2) The miss condition consisted of items that were not recognized during test (incorrect No response to test Question 1), In addition, several regressors were included to soak up BOLD variance associated with task aspects not included in any investigated contrast. During both encoding and retrieval, an item memory condition was included that consisted of items that were correctly recognized but for which the participant had no source memory (Yes response to Question 1 and No response to test Question 2 or incorrect response to Question 3) as well as a fourth regressor that modeled trials in which the participant did not produce any response to the first question. For the retrieval runs, four additional regressors were included to model the response to the new items (i.e. correct rejections and false alarms) and to model the second and third test questions (Questions 2 and 3). Temporal autocorrelations (AR(1)) in the residuals were corrected using a prewhitening approach. For difference in subsequent memory analyses (DSM; see below), a contrast of interest consisting of source - miss memory conditions was computed for each participant

### Hippocampal segmentation

Moving anteriorly through the coronal planes of an MNI-resampled human brain, y = −21 corresponds to the appearance of the uncus of the parahippocampal gyrus. In line with recent recommendations for long-axis segmentation of the hippocampus in human neuroimaging (Poppenk et al. 2013), we labeled hippocampal voxels at or anterior to this landmark as anterior HC while voxels posterior to the uncal apex were labeled as posterior HC. Specifically, for each participant, all functional voxels for which more than 50% of the underlying anatomical voxels were labeled as hippocampus by Freesurfer were considered functional representations of the hippocampus. While keeping the data in native subject space, we next established hippocampal voxels’ locations relative to MNI y = −21 by calculating the inverse of the MNI-transformation parameters for a given subject’s brain and projecting the back-transformed coronal plane corresponding to MNI y = -21 to functional native space. All reported activity- and connectivity measures thus represent averages from hippocampal subregions established in native space. A hippocampal segmentation from a representative subject is shown in Figure 1.

*Hippocampal connectivity. Correlational PPI analysis.* Task-dependent functional connectivity between hippocampal sub-regions was tested by correlational psychophysiological interaction (PPI) analyses (Fornito et al. 2012) implemented using routines from the gPPI toolbox for Matlab^®^ (McLaren et al. 2012). The PPI analysis allows studying connectivity shifts associated with task engagement while controlling for systematic variations in BOLD activity triggered by the experimental design. The correlational PPI analysis is a variation of the canonical PPI analysis in which the PPI term is derived from a partial correlation instead of a GLM, and thus creates symmetric PPI values between each pair of nodes. For each participant and HC sub-region (anterior, posterior × left, right), the time series from the first-level design matrix representing the different stimulus conditions of interest were multiplied separately by the deconvolved neural estimate and convolved with a canonical HRF, creating the PPI terms. The degree of connectivity between each pair of HC sub-regions was estimated trough partial correlations in which each HC subregion’s PPI time-series was partialled out by the original convolved task regressors and each HC sub-region’s time-series. Finally, all the PPI connectivity terms were z-transformed using Fisher’s r-to-z transformation. Source vs. baseline was used as the contras of interest for the PPI analyses.

### Statistical analyses

Generalized additive models (GAM) were run as implemented in the mgcv package for R (https://www.r-proiect.org) using Rstudio (www.rstudio.com) IDE. (Wood 2011, 2006). GAMs were run to test the continuous age-relationship of the different fMRI and task performance variables. For all fMRI analyses, estimated mean absolute and relative motion per participant were included as nuisance covariates. A smooth term for age was used. The smoothness of the age-curve is estimated as part of the model fit, and the resulting effective degrees of freedom (edf) was taken as a measure of deviation from linearity. The p-values associated with the smooth terms are only approximate, as they are based on the assumption that a penalized fit is equal to an unpenalized fit with the same edf, and do not take into account uncertainty associated with the smoothing parameter estimation. The major advantage of GAM in the present setting is that relationships of any degree of complexity can be modelled without specification of the basic shape of the relationship, and GAM is thus especially well-suited to map life-span trajectories of neurocognitive variables which can be assumed to be highly non-linear and where the basic form of the curve is not known (Fjell et al. 2010).

Separate GAMs were run for activity in aHC, pHC and the pHC-aHC difference, as well as the different memory performance variables (proportion of correctly remembered items with source memory [source memory hit], items with wrong source memory [wrong recollection], corrected source memory score [source memory hit - wrong recollection]). All activity analyses were run based on two different contrasts. First, activity during successful memory was compared to the implicit baseline. The results of this analysis reflected the difference in hippocampal activity during successful completion of the memory task compared to baseline activity, but did not reflect activity specifically associated with successful vs. non-successful memory. In the second set of analyses, difference in subsequent memory (DSM) was used as the measure of interest to allow isolation of activity related to memory success. [The term DSM is most often used to refer to encoding trials, but for simplicity we use the same term to refer to retrieval data also.] DSM was calculated as the difference in activity between items correctly recalled with source information (source trials) and activity to forgotten items (miss trials). To reduce the number of tests, these GAM analyses were run using mean values of right and left regions. Possible hemispheric differences were addressed by general linear model (GLMs) analyses especially suited to address such kind of interactions (see below).

To test for specific interaction effects, general linear models (GLM) with the factors HC axis (anterior, posterior) × condition (encoding, retrieval) × hemisphere (right, left) × accuracy (source-baseline, source-miss) × age group (8 groups) × Sex (female, male), with motion (mean absolute and relative) as nuisance variables. The use of 8 age groups was motivated by the results of the preceding GAM fits which showed that the age-trajectories were too complex (edf > 6) to be modelled with fewer age groups.

Further, we tested whether fMRI activity was related to memory performance by running additional GAMs with corrected source memory as dependent variable, and age and brain activity as smooth terms. This was done for aHC, pHC as well as the pHC-aHC difference, for both the source memory - baseline contrast and the DSM effect.

In the next set of analyses, we tested the lifespan trajectories of functional connectivity between aHC and pHC, both within and across hemispheres. We estimated functional connectivity within the PPI framework. In short, we tested whether connectivity between hippocampal regions were different during the memory task than during baseline. This difference was then quantified and used in further GAM and GLM analyses. All the higher-level statistical analyses (but GAM fitting) were run in SPSS (version 24).

## Results

### Behavioral results

Mean corrected source memory score was 0.44 (SD = 0.18). Scatterplots illustrating individual scores in memory performance against age are shown in Figure 2. GAMs were run with the corrected source memory score, source memory hits and wrong recollection in turn as independent variables and age as smooth term. Age was in all cases highly significantly related to memory (all p’s < 2e^-16^), and the trajectories were also highly non-linear (corrected source memory score adjusted R^2^ = .37, edf = 5.25; source memory hits adjusted R^2^ = .18, edf = 5.69; wrong recollection adjusted R^2^ = .31 edf = 3.87). The mean number of trials included for correct source memory and miss memory was 51.60 (Sd = 10.60) and 23.05 (SD = 14.39), respectively. Detailed descriptives for all response classes are presented in Supplemental information.

**Figure 2.**
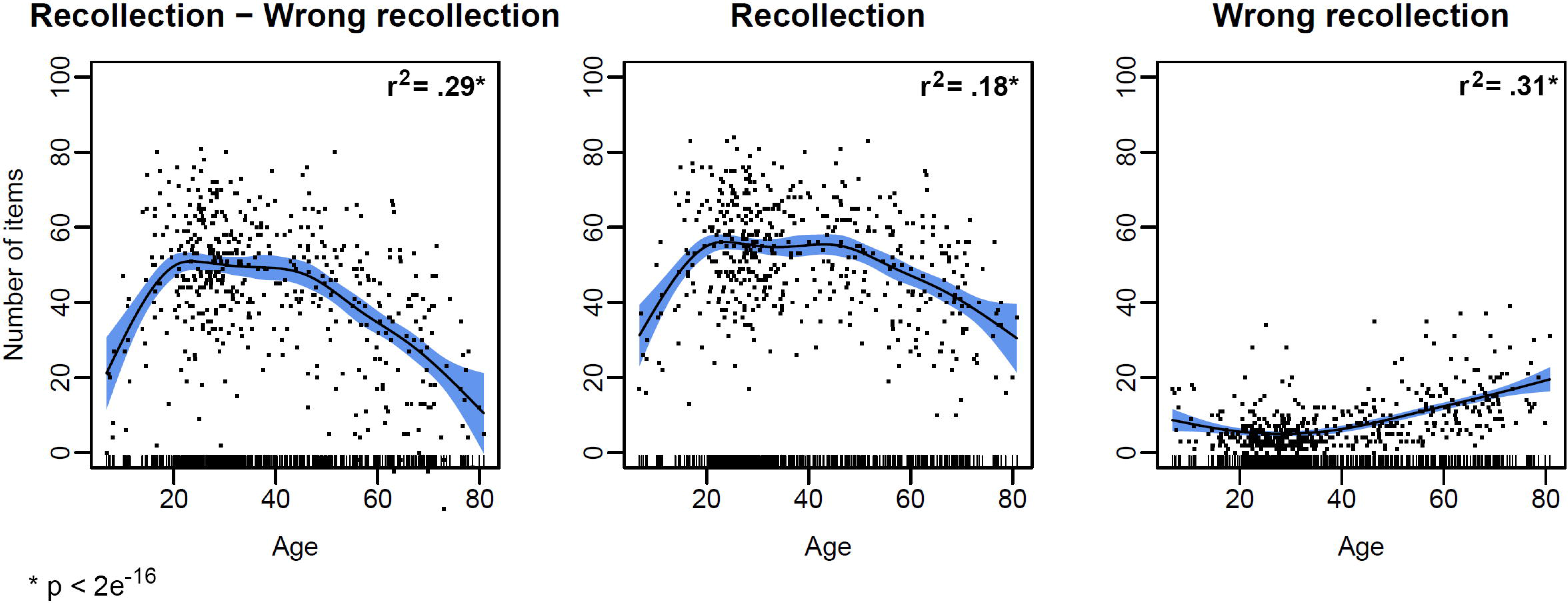
Lifespan trajectories of performance. Individual memory performance scores plotted across age. The curves represent the smooth function of age from the generalized additive models. The left panel represent the corrected source memory score (recollection - wrong recollection) used in all analyses. The shaded area around the curves represent 2 standard errors of the mean. * p < .05.

### Hippocampal activity - age trajectories

First, to map the general effects of age on activation in anterior and posterior hippocampus, GAM models with activity as dependent variables and age as a smooth predictor were run, with movement as nuisance covariates. Scatterplots with imposed GAM fits corrected for motion parameters are shown in Figure 3. For encoding, no significant effects of age were found for either anterior (Baseline contrast: edf = 1.0, F = 0.8, adj R^2^ = 0, p = .4/ DSM: edf = 1.7, F = 0.7, adj R^2^ = 0, p = .5) or posterior (Baseline contrast: edf = 1.0, F = 2.4, adj R^2^ = 0.01, p = .12/ DSM: edf = 1.2, F = .6, adj R^2^ = 0, p = .6) hippocampal activity. However, a significant negative effect of age on the posterior-anterior source memory vs. baseline contrast (Baseline contrast: edf = 1, F = 6.6, adj R^2^ = .02, p = .011/ DSM: edf = 1.0, F = 0.2, adj R^2^ = 0, p = .67) was found.

**Figure 3.**
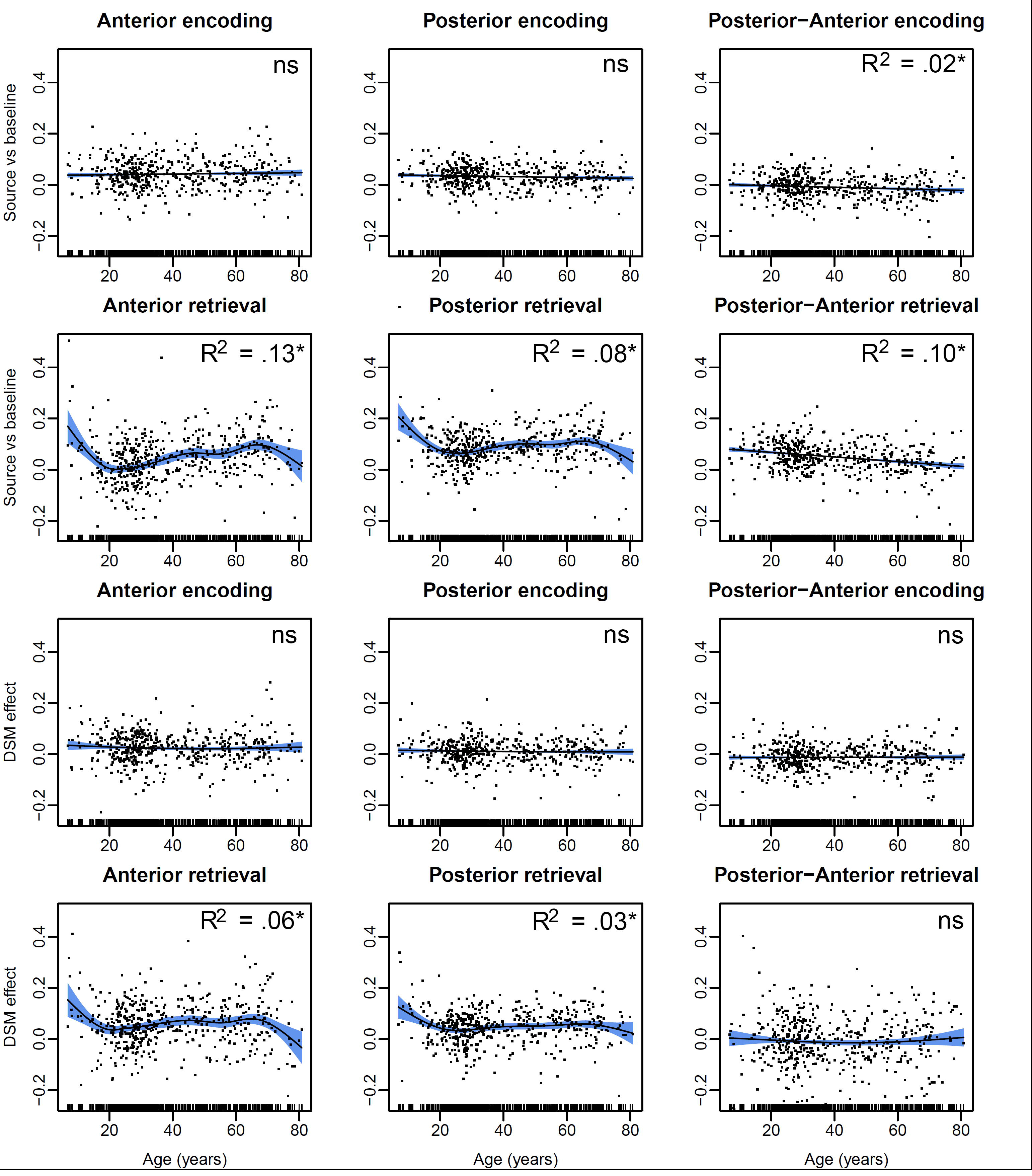
Lifespan trajectories of fMRI activity. Individual fMRI contrast values plotted across age. The curves represent the smooth function of age from the generalized additive models. The plots are residualized on absolute and relative movement during scanning. The two top rows depict values based on the difference between source memory and implicit baseline. The two bottom rows depict the difference in subsequent memory (DSM) effect, i.e. the difference in activity to correct source memory vs. miss trials. “Posterior-Anterior” represents the difference between posterior and anterior hippocampal activity. The shaded area around the curves represent 2 standard errors of the mean. * p < .05, ns: not significant.

For retrieval, however, robust age effects were seen for both anterior (Baseline contrast: edf = 6.4, F = 9.3, adj R^2^ = .13, p = 4.61e^−11^ / DSM: edf = 5.6, F = 3.7, adj R^2^ = 0.06, p = .0007) and posterior (Baseline contrast: edf = 6.0, F = 6.4, adj R^2^ = .08, p = 3.01e^−07^/ DSM: edf = 5.0, F = 3.2, adj R^2^ = 0.03, p = .004) hippocampal activity. Inspections of the trajectories showed a highly non-linear pattern across the life-span, which was confirmed by the high edf values. Both the anterior and the posterior source memory vs. baseline effects and the DSM effects were smaller with advancing age during development From about 20 years, however, the curves were positive, indicating higher activity with higher age, before a negative slope was observed from about 70 years. This pattern was somewhat more evident for the anterior compared to the posterior hippocampus.

Interestingly, at retrieval, a direct test of the posterior-anterior DSM effects did not show a significant effect of age for DSM (edf = 1.8, F = 0.6, R^2^ = 0, p = .26) but a significant effect for the baseline contrast (edf = 1, F = 42, R^2^ = 0.10, p = 2.12e^-10^). The posterior advantage was high in early development, and showed a linear negative relationship with age throughout the age span. The lack of effects when using the DSM contrast suggests that the age effect on the anterior-posterior differentiation is related to execution of the retrieval task per se, not the successful retrieval of episodic memory content.

### Hippocampal activity - interaction effects

A GLM with the factors long-axis (anterior, posterior) × condition (encoding, retrieval) × hemisphere (right, left) × accuracy (source-baseline, miss-baseline) × age group (8 groups) × sex (female, male) were run. The results are presented in Table 2, and only the most relevant results will be highlighted here.

**Table 2.** General linear model results. Age was divided in 8 different groups and entered as a factor in the analysis. The Greenhouse-Geisser method was used for correction of violation of sphericity. Absolute and relative movement and sex were used as covariates.
Contrasts of no interest were omitted from the table Accuracy refers to the source memory - baseline contrast vs. the miss - baseline contrast. Condition refers to encoding vs. retrieval. Hemi refers to hemisphere. **Bold** indicates p < .05

|  | F | p < |
| --- | --- | --- |
| *Within-subject effects* |  |  |
| Accuracy | 47.30 | **1.9e-11** |
| Accuracy × Age | 4.13 | **.001** |
| Accuracy × Condition | 11.17 | **.001** |
| Accuracy × Age × Condition | 2.69 | **.01** |
| Accuracy × Anterior-Posterior | 0.01 | .91 |
| Accuracy × Anterior-Posterior × Age | 0.64 | .72 |
| Accuracy × Anterior-Posterior × Condition | 5.81 | **.05** |
| Anterior-Posterior | 6.58 | **.05** |
| Anterior-Posterior × Age | 8.32 | **1.3e-9** |
| Hemi | 1.48 | .23 |
| Hemi × Age | 0.85 | .55 |
| Condition | 1.39 | .24 |
| Condition × Age | 4.40 | **.0001** |
| Anterior-Posterior × Hemi | 0.13 | .71 |
| Anterior-Posterior × Hemi × Age | 0.31 | .95 |
| Anterior-Posterior × Condition | 10.62 | **.001** |
| Anterior-Posterior × Condition × Age | 1.80 | .09 |
| Hemi × Condition | 2.62 | .11 |
| Hemi × Condition × Age | 0.92 | .49 |
| Anterior-Posterior × Hemi × Condition | 0.69 | .41 |
| Anterior-Posterior × Hemi × Condition × Age | 0.74 | .64 |
| *Between-subject effects* |  |  |
| Age | 3.89 | **.001** |

There was a strong effect of accuracy, with higher activity for source vs. baseline than miss vs. baseline (see Figure 4). This confirms that the paradigm produced the expected DSM effect. There was an accuracy × age interaction, with children showing larger difference between activity for source memory vs. miss. Also, we found an anterior-posterior activity × condition interaction, with aHC showing higher activity during encoding than during retrieval, and the opposite pattern for the pHC. There was a strong age × anterior-posterior interaction, caused by generally higher activity in the posterior than the anterior hippocampus in children and young adults, with the difference gradually diminishing in older adulthood. Finally, there was a tendency (p = .09) towards an age × anterior-posterior × condition interaction. This appeared due to lower anterior-posterior activity differentiation during retrieval in older age, combined with a tendency for higher anterior than posterior encoding activity in middle adulthood.

**Figure 4.**
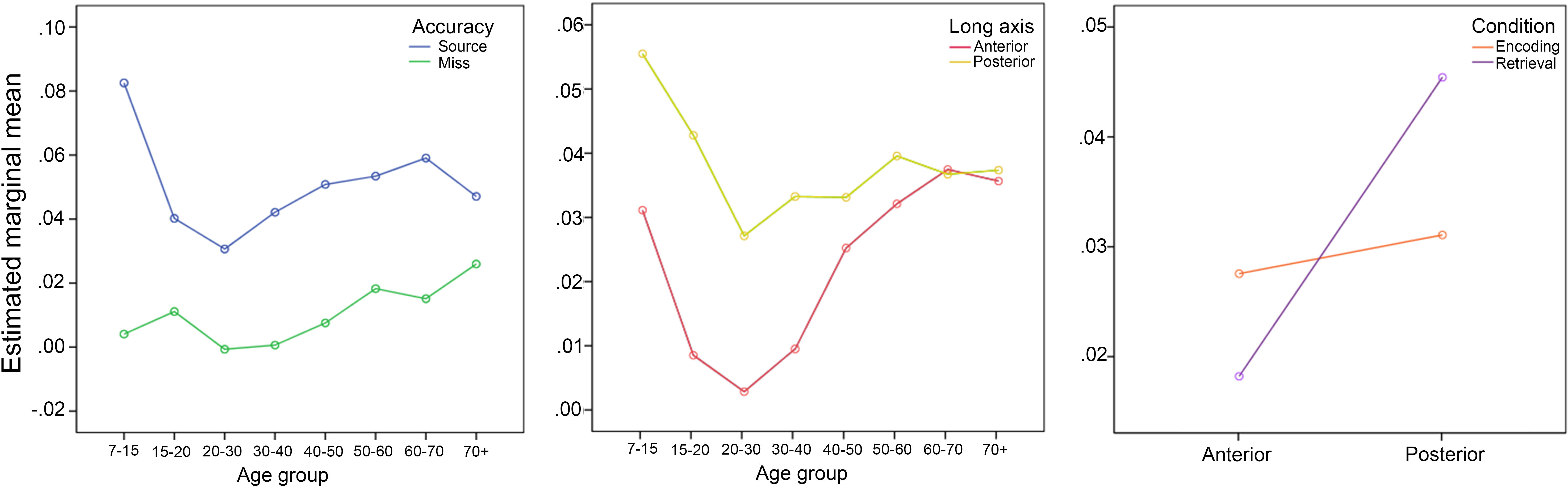
Interaction plots. Left panel: Activity associated with source memory vs. miss across age groups. Middle panel: Source memory-related activity in anterior and posterior hippocampus across age groups (baseline contrast). Right panel: Evidence for the long-axis anterior-posterior encoding-retrieval specialization (Source memory-related activity, baseline contrast). The plots are residualized on sex and movement during scanning. The interactions represented by each plot were all significant (p < .05).

These results showed the expected hippocampal long axis specialization for encoding vs. retrieval, and that the long-axis activity was related to age. The lack of a significant condition × anterior-posterior interaction means that the hippocampal long axis effects did not reflect activity related to successful vs. unsuccessful [miss] source memory, but rather general activity during performance of memory tasks. However, there was also a condition × anterior-posterior × condition interaction, which was caused by a breakdown of the expected long-axis encoding-retrieval specialization during encoding miss trials (see Figure 5).

**Figure 5.**
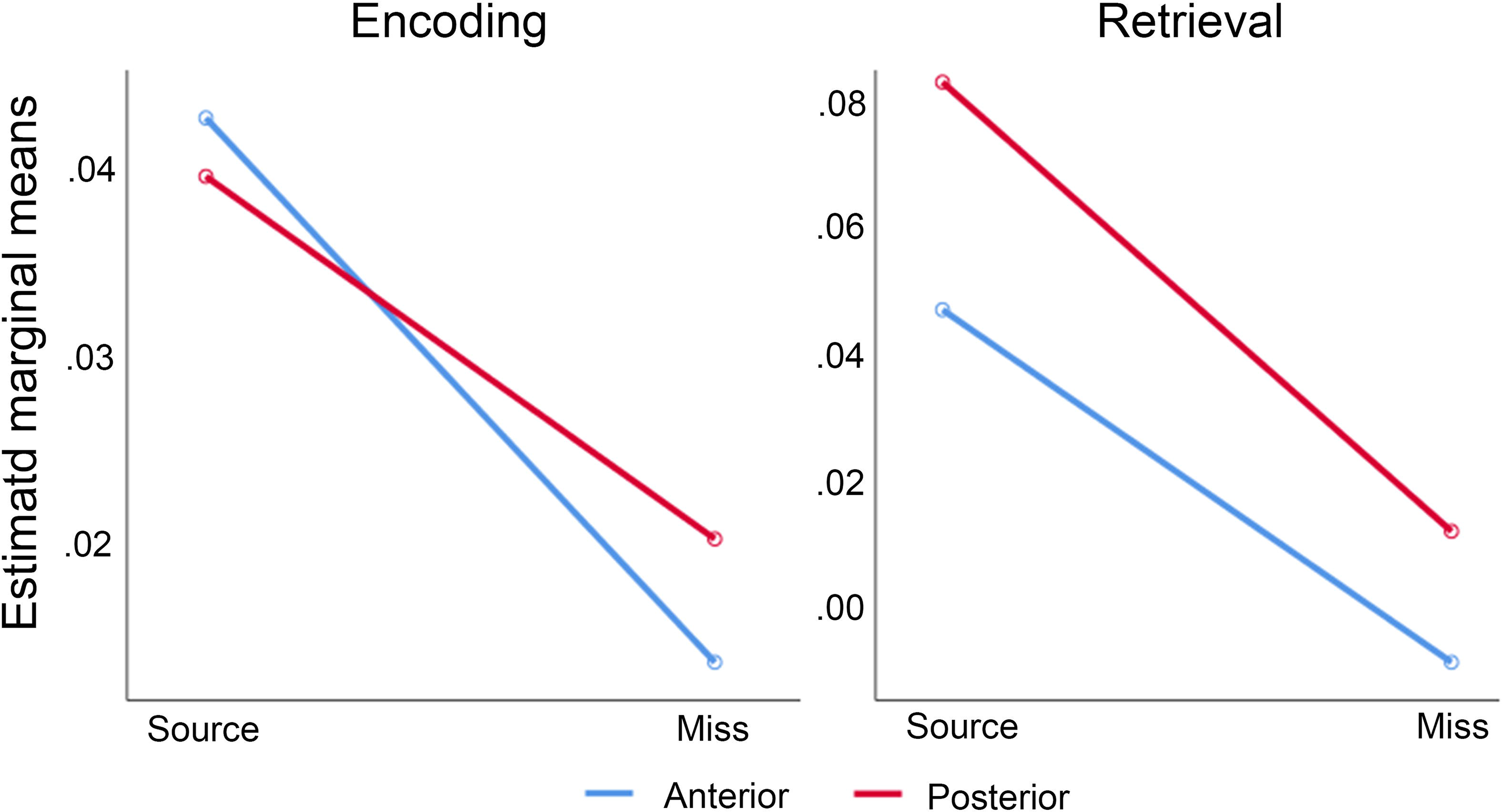
Breakdown of specialization during unsuccessful memory. There was a significant accuracy × anterior-posterior × condition interaction, which was caused by the long-axis encoding-retrieval specialization to break down during encoding miss trials.

### Hippocampal activity - relationships with performance

GAM models were run with the corrected source memory score as dependent variable, and age and hippocampal activity as smooth terms, with the movement variables as covariates. Separate models for aHC and pHC, and for encoding and retrieval were tested. Additional models included the posterior-anterior difference as predictor. The analyses were run both for the baseline contrast and for the DSM contrast. The relationships were weak, with only one reaching an uncorrected significance level of p < .05 (aHC retrieval, *t* = 2.12, p = .035). Due to the number of tests, this relationship did not survive proper correction for multiple comparisons (False Discovery Rate [FDR]) and was thus not considered further. Since these analyses did not reveal other significant relationships, further analyses testing interactions were not performed.

### Functional connectivity - psychophysiological interaction analysis

The age-trajectories for task-related hippocampal connectivity are shown in Figure 6. Connectivity between all hippocampal regions, i.e. aHC vs pHC and right vs left hemisphere, was higher during task than during baseline (all p’s <le^−16^, see Table 3). The degree of connectivity between hippocampal regions was similar during encoding and retrieval (F = .8, p = .4, see Table 4). A main effect of connectivity region (F = 255.3, p < 1e^−16^) revealed that the highest task-related connectivity was observed for aHC - pHC connectivity within the same hippocampus, both for the left and the right hemisphere. Further, inter-hemispheric connectivity was higher for aHC than pHC during the memory task. Both these effects were observed regardless of condition, i.e. encoding versus retrieval.

**Figure 6.**
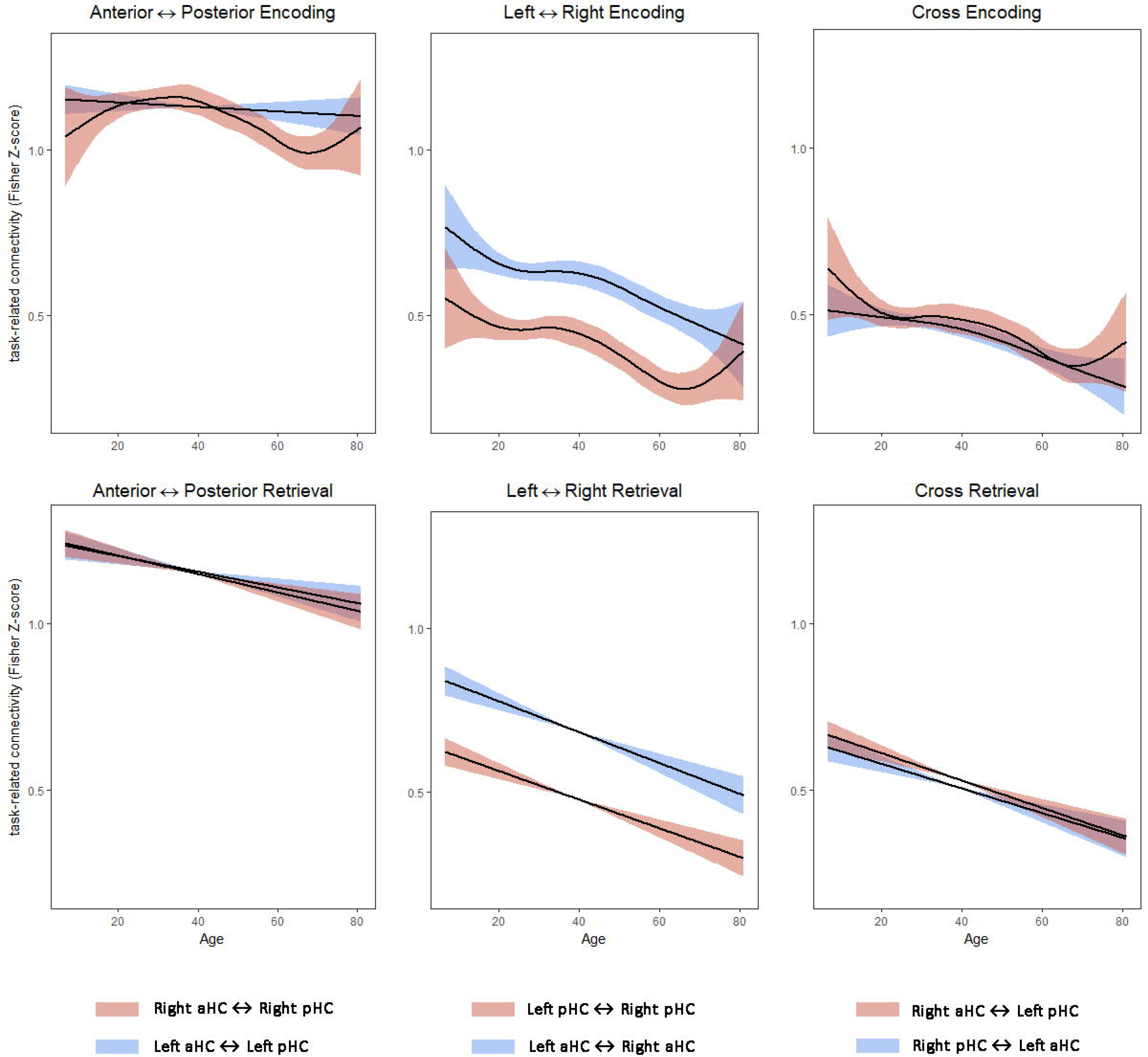
Lifespan trajectories of task-related connectivity. The curves represent the smooth function of age from the generalized additive models. The fit line is residualized on absolute and relative movement during scanning. The shaded area represents 2 standard errors of the mean.

**Table 3.** Task (source vs. baseline) related changes in connectivity. Statistics represent one-sample t-test with significance adjusted for multiple comparisons with FDR (n = 12).

| PPI - Connectivity pair |  | Task - Baseline |  |
| --- | --- | --- | --- |
|  |  | *t* | *p* |
| ENCODING | Left aHC – Left pHC | 101.06 | <1e<sup>-16</sup> |
|  | Left aHC – Right aHC | 63.69 | <1e<sup>-16</sup> |
|  | Left aHC – Right pHC | 50.12 | <1e<sup>-16</sup> |
|  | Left pHC – Right aHC | 50.60 | <1e<sup>-16</sup> |
|  | Left pHC – Right pHC | 46.15 | <1e<sup>-16</sup> |
|  | Right aHC – Right pHC | 113.91 | <1e<sup>-16</sup> |
| RETRIEVAL | Left aHC – Left pHC | 108.72 | <1e<sup>-16</sup> |
|  | Left aHC – Right aHC | 63.72 | <1e<sup>-16</sup> |
|  | Left aHC – Right pHC | 53.92 | <1e<sup>-16</sup> |
|  | Left pHC – Right aHC | 55.90 | <1e<sup>-16</sup> |
|  | Left pHC – Right pHC | 51.58 | <1e<sup>-16</sup> |
|  | Right aHC – Right pHC | 109.94 | <1e<sup>-16</sup> |

**Table 4.**
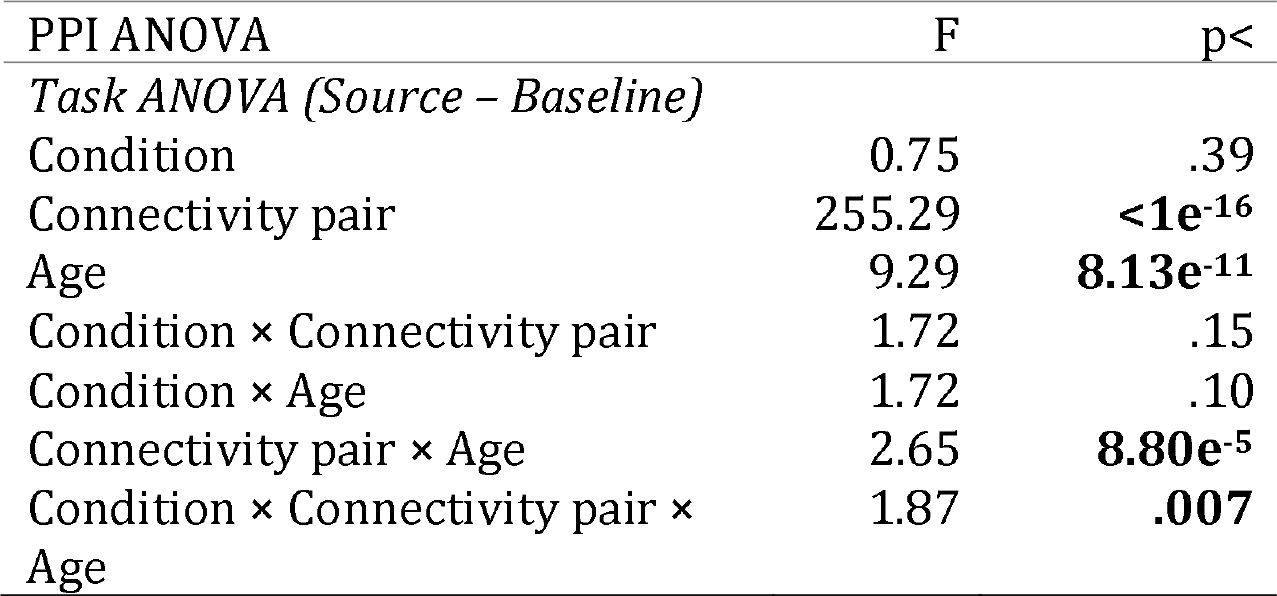
General linear model results for PPI. Age was divided in 8 different groups and entered as a factor in the analysis. The Greenhouse-Geisser method was used for correction of violation of sphericity. Absolute and relative movement and sex were used as covariates.
The F and *p* statistics represent results from two GLMs where PPI coefficients from source memory vs. baseline were introduced as the predicted variable. Contrasts of no interest were omitted from the table **Bold** indicates p < .05

In general, task-related connectivity between sub-regions showed a monotonic reduction with increasing age (main effect of age [F = 9.3, p = 8.1e^−11^]). For most connections, the youngest children showed the highest connectivity which then decreased continuously throughout adulthood. This general trend was consistent both for encoding and retrieval connection, except for intra-hippocampal anterior-posterior encoding connectivity which remained somewhat stable across life. This exception was reflected in a three-way interaction between age × connectivity pair × condition (F = 1.9, p = .007).

Further, we tested whether task-related connectivity correlated with memory performance. No significant relationships were observed independently of age, neither during encoding nor during retrieval (all tests p > .05 uncorrected).

## Discussion

Three main sets of findings were obtained: First, we found the encoding-retrieval specialization along the hippocampal long axis, with higher anterior activity during encoding and higher posterior activity during retrieval. The anterior-posterior differentiation was more pronounced during retrieval. While this was according to our hypothesis, the novel result was that long-axis specialization was seen during engagement in an encoding and retrieval task, not depending on the successful completion of this task. Thus, the long-axis hippocampal specialization characterized modi operandi for encoding and retrieval, respectively, more than being specific to successful source memory processes. Second, degree of long-axis differentiation correlated with age, mainly caused by higher posterior activity in children, with gradually higher anterior activity in adulthood and older age. Age effects on long-axis differentiation was according to our hypothesis, but we expected an inverse U-shaped rather than a linear age-relationship. Thus, long-axis differentiation across the lifespan did not adhere to a “from-less-specialized to more-specialized” principle in development and the inverse in aging. Further, while retrieval activity showed a markedly non-linear age-trajectory, encoding activity was stable across age. The retrieval trajectory indicated a rapid developmental phase and substantial reduction in aging. Finally, FC between aHC and pHC increased during the memory task, and the degree of increase was related to age for all within- and between hippocampus connections for both encoding and retrieval, except for the anterior-posterior during encoding. As expected, children showed unspecific task-related increase in FC both within and between hemispheres, possibly indicating that their ability to distribute tasks to more specialized regions is not yet mature. However, we did not observe increased connectivity among the older adults, as we would expect if lower connectivity indexes higher degree of specialization. The implications of these main findings are discussed below.

### Long axis specialization for encoding vs. retrieval

Support for the long axis specialization for encoding an retrieval in humans comes from functional brain imaging studies (Lepage, Habib, and Tulving 1998), including meta-analyses (Spaniol et al. 2009; Kühn and Gallinat 2014). The present results quite clearly confirmed this pattern. There was no sign of an interaction between accuracy and anterior-posterior axis, showing that it did not matter for the long-axis specialization whether an item eventually was remembered with source or forgotten. The authors of a previous meta-analysis suggested that the specific contrast used to define encoding and retrieval success was important, since different contrasts reflect activity differences between cognitive processes that activate hippocampal sub-regions to various degrees (Spaniol et al. 2009). The present results suggest that the hippocampal long-axis specialization for encoding vs. retrieval exists, but that it reflects engagement in the memory task, not the successful encoding or retrieval of episodic memories. This specialization was mostly driven by retrieval, where activity was much higher in pHC vs. aHC. During encoding, aHC and pHC activation was comparable. Nevertheless, the evident differences between retrieval vs. encoding activity along the long axis justified further testing age effects on hippocampal regional differentiation.

### Age effects on hippocampal long axis differentiation

We made two major observations. First, the hippocampal long-axis differentiation was linearly negatively related to age throughout the age-span of almost 75 years. Thus, we did not identify a clear developmental end-point or aging-related start-point. Rather, the posterior preference in the youngest children was replaced by gradually higher aHC relative to pHC activity with higher age, continuing for most of the life-span. This indicates that the hippocampal long-axis differentiation is sensitive to age both in development and aging, but not in a simple “less-specialization-versus-more-specialization” framework. The results fit better with a posterior-to-anterior shift in activity, suggested to characterize the development to adulthood phase (Sastre Iii et al. 2016). Some of these effects may be related to ongoing structural maturation and age-reductions of the hippocampus (Ostby et al. 2009; Krogsrud et al. 2014; Daugherty et al. 2016; Walhovd et al. 2005), which has been related to episodic memory performance (Ostby et al. 2012; Tamnes et al. 2014; Lee, Ekstrom, and Ghetti 2014; Keresztes et al. 2017; Daugherty, Flinn, and Ofen 2017; DeMaster et al. 2014; Fjell et al. 2013). These studies suggested that differential maturation of hippocampal subfields is relevant for development of episodic memory, which fits with the present fMRI results.

Importantly, the age effect on the hippocampal long-axis differentiation was not seen for the DSM effect, suggesting that this effect is not specific to successful memory and hence does not necessarily reflect successful retrieval of episodic memory content. This is in line with a previous aging study finding long-axis hippocampal specialization of anterior-encoding and posterior-retrieval by use of a non-memory contrast task (Salami, Eriksson, and Nyberg 2012).

The second major finding was that retrieval activity was much more sensitive to age than encoding activity, both in development and aging. Even though we found a significant effect of age on long-axis differentiation also for encoding, encoding activity was not related to age in aHC or pHC per se. In contrast, retrieval activity showed complex age-functions. Highest activity was seen in the children, with a negative trajectory suggesting a developmental end-point at around 25 years. From this age, a positive age-relationship was seen up to almost 70 years, after which the curve again was negative towards the end of the age-range. The observation that activity during retrieval in older adults approached the same levels as that seen in children is interesting, and could reflect more effortful processing. ‘Over-recruitment’ in aging has for instance been interpreted as a sign of neural inefficiency (Duverne, Habibi, and Rugg 2008). The high levels of activation are not likely to be a direct result of higher task demands in these groups or reflect task performance per se, as activity was not correlated with memory performance, and the age-effects were clearly attenuated when the DSM effect was used instead of the baseline contrast The gradual increase in anterior relative to posterior hippocampal retrieval activity through development and into young adulthood fits well with the results of a previous large study (n = 126) (Sastre Iii et al. 2016). In that study, age-differences occurring along the longitudinal axis were identified, with selective activation in the hippocampal head in high performing adults but not in children.

The presently observed age-effects are in line with some previous fMRI studies (DeMaster and Ghetti 2013; Paz-Alonso et al. 2008) but not others (Guler and Thomas 2013; Ofen et al. 2012; Ofen et al. 2007), which reflects that age effects on hippocampal memory-related activity are not universally found. The combination of a continuous and wide age-range combined with scanning during both encoding and retrieval, and the use of two different contrasts to define memory activity, enables us to shed some light on the conditions for finding age effects on hippocampal activation. First, age effects were much larger for retrieval than encoding activity both in development and aging. Second, age effects were clearly attenuated when a DSM contrast was used instead of a baseline contrast, which may indicate that hippocampal activity related to memory success per se (Cansino et al. 2015) may be less sensitive to age than activity related to performance of a memory task not specifically depending on memory success (Salami, Eriksson, and Nyberg 2012). Third, retrieval activity showed a complex, non-monotonous trajectory, which means that continuous sampling across larger age ranges will likely yield more information than comparing groups of restricted age. Finally, although age effects tended to be stronger in aHC than pHC, implying that studying sub-regions may increase age-sensitivity (Carr et al. 2017; Sastre Iii et al. 2016), this factor was of relatively less importance in the present study than those discussed above. To conclude, even though these factors cannot explain all differences in reported hippocampal activity in previous development and aging studies, we suggest that age most strongly correlate with anterior, retrieval-related differences between source memory and baseline activity, and that posterior, encoding-related differences between successful vs. unsuccessful source memory activity will be relatively more resistant to the influence of age.

### Age effects on task-related connectivity

Psychophysiological interaction analysis has been applied in studies of development and aging of episodic memory. Development of connectivity between medial temporal lobe and prefrontal cortical regions during retrieval appears to continue into young adulthood (Ofen et al. 2012). In a study of adults, FC increases were generally reduced with higher age, but such a reduction did not apply to hippocampal connections (King, de Chastelaine, and Rugg 2017). Age effects on intra-hippocampal connectivity-changes during memory tasks have to our knowledge not been tested. As hypothesized, we found increased FC within and between hippocampi during encoding and retrieval. This increase was much higher within the hippocampus than between homologous contralateral regions. Of most relevance for the present study, the children showed higher connectivity increases than any other age group. This was seen for all tested connections except anterior-posterior FC during encoding. The high FC in the children could in principle signify less developed sub-regional specialization of communication, less efficient neural processing or lack of inhibition. However, the mostly linear age-trajectories indicate that if high connectivity in the children signify lack of maturation, then the older adults may appear to have the most efficient FC, which is a less likely interpretation. As we did not find a correlation between FC and memory performance, the findings are more in line with the children using their brains differently than adults to accomplish the task. More research is needed to understand the implication of the age effects on hippocampal FC. Still, these results indicate that in addition to age-effects on hippocampal sub-region activity, there are also substantial differences in task-related hippocampal connectivity across the lifespan when participants are engaged in memory encoding and retrieval tasks.

### Limitations

There are multiple limitations with the present study. First, since all analyses are based on cross-sectional analyses, the results only regards age differences, not changes per se. Few longitudinal fMRI studies of development or aging exist (for exceptions, see (Pudas et al. 2013; Persson et al. 2012)), but cross-sectional analyses can sometimes yield spurious results, at least in aging (Nyberg 2017). Further, although the division of hippocampus in an anterior and a posterior section has merit (Poppenk et al. 2013), this division crosses established hippocampal subfields running along the anterior-posterior axis (Fanselow and Dong 2010). With higher field strengths, e.g. 7T, other subfield divisions could be valuable to test (Carr et al. 2017; Berron et al. 2016). Further, as discussed above, a long-axis specialization may be valid for many types of cognitive processes (Poppenk et al. 2013; Maass et al. 2014; Kim 2015; Moscovitch et al. 2016), which means that subtle differences in task demands may alter the relative impact of aHC vs pHC, and possibly also their lifespan trajectories. Finally, movement during scanning may affect the results, especially when children and older adults are included. In the present study, care was taken to minimize movement during scanning, to remove effects of movement during pre-processing, and both absolute and relative movement were included as covariates in the analyses.

### Conclusion

We identified a clear anterior-posterior hippocampal long-axis specialization for encoding vs. retrieval, which was linearly related to age across almost 75 years. Still, a posterior-to-anterior shift in activity from childhood to adulthood was the most prominent age-effect on the hippocampal long-axis. This effect was driven by the strong influence of age on retrieval activity both in development and aging. Connectivity between hippocampal sub-regions increased during execution of the memory task, and children showed substantially higher connectivity than adults. Effects were seen when source memory activity was compared to baseline activity, irrespective of accuracy. Thus, the hippocampal long-axis specialization and differentiation seemed to reflect task engagement more than processes relayed to successful performance of the task.

## Acknowledgement

We thank Lars Nyberg for valuable comments to an earlier version of the manuscript.

